# Assembly of the amphibian microbiome is influenced by the effects of land-use change on environmental reservoirs

**DOI:** 10.1101/2020.11.30.405050

**Authors:** Elle M. Barnes, Steve Kutos, Nina Naghshineh, Marissa Mesko, Qing You, J.D. Lewis

## Abstract

A growing focus in microbial ecology is understanding of how beneficial microbiome function is created and maintained through both stochastic and deterministic assembly mechanisms. This study explores the role of both the environment and disease in regulating the composition of microbial species pools in the soil and local communities of an amphibian host. To address this, we compared the microbiomes of over 200 *Plethodon cinereus* salamanders along a 65km land-use gradient in the greater New York metropolitan area and paired these with associated soil cores. Additionally, we characterized the diversity of bacterial and fungal symbionts that putatively inhibit the pathogenic fungus *Batrachochytrium dendrobatidis*. We predicted that if soil functions as the main regional species pool to amphibian skin, variation in skin microbial community composition would correlate with changes seen in soil. We found that salamanders share many microbial taxa with their soil environment but that these two microbiomes exhibit key differences, especially in the relative abundances of the bacteria phyla Acidobacteria, Actinobacteria, and Proteobacteria and the fungal phyla Ascomycota and genus *Basidiobolus*. Microbial community composition varied with changes in land-use associated factors such as canopy cover, impervious surface, and concentrations of the soil elements Al, Ni, and Hg, creating site-specific compositions. In addition, high dissimilarity among individual amphibian microbiomes across and within sites suggest that both stochastic and deterministic mechanisms guide assembly of microbes onto amphibian skin, with likely consequences in disease preventative function.

## 1 Introduction

Soil microorganisms perform a number of key functions in forest ecosystems. Although they are most well known for their roles in organic matter decomposition and nutrient cycling, they also contribute to the microbial diversity of a number of terrestrial host organisms, including both plants (Bakker et al., 2014; Chaparro et al., 2012; Mendes et al., 2013; Smith & Read, 2010; van der Heijden et al., 1998) and animals (Bahrndorff et al., 2016; Ross et al., 2019). For example, soil can act as a reservoir, or supplier, of microbial taxa to many amphibian hosts (Loudon et al., 2014; Muletz et al., 2012). Thus, even if anthropogenic activities do not appear to directly impact amphibians, changes in the soil microbial reservoir may negatively influence their microbiome (Bird et al., 2018; Ficetola et al., 2015; Varela et al., 2018). Despite the implication of soil-obtained microbes in a number of important processes such as disease protection and skin functionality (Becker & Harris, 2010), the indirect role that environmental-driven changes in the soil reservoir play in amphibian host health is understudied. In situations where the amphibian microbiome is disturbed (e.g. during *Batrachochytrium dendrobatidis* (Bd) infection), recolonization of microbiota onto a host is likely dependent on not only the composition of the reservoir and host identity, but also on stochastic assembly mechanisms leading to distinct microbial communities across individuals (Callens et al., 2018; Loudon et al., 2014; Muletz-Wolz et al., 2018).

For more than half a century, ecologists have used the process of community assembly to describe the formation of species diversity from regional species pools to local communities influenced by both contemporary and historical factors (Elton, 1946; Hubbell, 2005; Leibold et al., 2004; MacArthur & Wilson, 2001; Patrick, 1967; Williams,1947). Contemporary factors, such as the abiotic and biotic conditions of a local community, can drive particular species distributions in a deterministic manner based on species’ niche requirements (Baas-Becking, 1934; Cottenie, 2005; Logue et al., 2011). Alternatively, historical factors, such as dispersal limitation, drift, and priority effects, can be driven by past stochastic events that have lasting effects on current species distributions (Andersson et al., 2014; Fukami, 2015; Hanson et al., 2012; Leibold et al., 2010). These two factors have been explored and identified in many systems: plants, animals, and more recently in microbes.

A growing body of evidence suggests that both free-living and host-associated microbes fluctuate in abundance, distribution, and composition across both space and time (Fierer & Jackson, 2006; Martiny et al., 2006; Papke & Ward, 2004; Peay et al., 2016; Tedersoo et al., 2012, 2014). For instance, studies have found that soil microbes are sensitive to local moisture conditions, temperature, pH, and nutrient concentrations and availability, suggesting that microbial biogeography is not entirely random (Fierer & Jackson, 2006; Hansel et al., 2008; Treseder, 2004; Tedersoo et al., 2020; Vartoukian et al., 2010). Regional environmental gradients, such as those across land-use change, have enabled researchers to explore microbial biogeography over space while also reducing confounding effects of broad continental climatic factors (Arnfield, 2003; Epp Schmidt et al., 2017; Fierer et al., 2011; Grimm et al., 2008; Looby et al., 2016; McDonnell et al., 1997; McDonnell & Pickett, 1990; Peay et al., 2017; Pouyat et al., 1995). The results of these studies suggest that environmental variation over even small geographic scales can directly affect soil microbial community composition, abundance, and functional diversity (Carrino-Kyker et al., 2011; Charlop-Powers et al., 2016; Lilleskov et al., 2002; Reese et al., 2016; Xu et al., 2014; Yan et al., 2016).

In this study, we explored the bacterial and fungal diversity of *Plethodon cinereus* salamanders and their soil reservoir along a gradient of land-use change in New York, USA. For Plethodontid salamanders, habitat conditions in the local environment appear to be key drivers of cutaneous microbial diversity (Bird et al., 2018; Varela et al., 2018). Given this, we predicted that 1) land-use change, from exurban to urban habitats, would impact local soil microbial community structure due to changes in soil physicochemical characteristics such as pH and soil nutrient concentrations (De Carvalho et al., 2016; Ge et al., 2008), and 2) changes in the soil microbial community would ultimately impact which microbial species were available to colonize salamander skin (Bletz et al., 2016). Additionally, we predicted that soil and salamander microbiomes would be distinct from one another with salamander skin being less diverse than their soil environment and with differences in the relative abundance of dominant microbial taxa (Kueneman et al., 2019). Finally, we predicted that there would be an increased abundance of anti-Bd microbial taxa associated with salamander skin as opposed to the soil environment due to increased pathogen loads on hosts (Barnes et al., 2020; Loudon et al., 2016; Rebollar et al., 2016). Overall, our study offers important new insights into the indirect effects of land-use change on host-associated microbiota as well as identifies key factors guiding assembly, diversity, and function of the amphibian microbiome.

## 2 Methods

### 2.1 Field Sampling

We collected skin swabs from the terrestrial woodland salamander species, *Plethodon cinereus* (*n* = 211), and soil cores (*n* = 94) from nine sites along a 65-km gradient of land-use change in Spring and Fall of 2017-2018 (**Table S1**). Of these nine sites, three were categorized as urban (Van Cortlandt Park (VC), New York Botanical Gardens (BG), and Pelham Bay Park(PB)), three as suburban (Rockefeller State Park Preserve (RF), Fordham University Louis Calder Center (CA), and Westmoreland Nature Sanctuary (WM)), and three exurban (Hudson Highlands State Park (HH) and two sites within Fahnestock State Park (FS1 & FS2)), based on percent impervious surface, population density, and distance from Central Park, NY, USA (Dewitz, 2019; New York State Department of Health). These sites were chosen because they are east of the Hudson River, have acidic loamy soils, and share similar bedrock parent material, forest stand age and diversity (Brock & Brock, 2001; Edinger, 2014; McDonnell et al., 1997; Schuberth, 1968; Soil Survey Staff, 1975; Thornes, 1974). This allowed us to minimize confounding variables associated with the regional land-use gradient. Past studies of the region have shown many impacts to local soils from anthropogenic land-use in sites classified as urban and suburban (Lewis, 2013; Pouyat & McDonnell, 1991; Pouyat et al., 1995; Pouyat et al., 2008; Schuler, 2011; White & McDonnell, 1988).

A non-invasive swabbing method was used based on Boyle et al. (2004). Salamanders were bathed in sterile water to remove transient microbes and then swabbed using a sterile cotton swab on their ventral, right, and left sides (Harris et al., 2006). Three replicate soil cores were taken in a 0.25 m-radius around salamander sampling locations and < 1 meter from the bole of a mature *Fagus grandifolia* (DBH > 30 cm) located in each sampling plot. Soil cores were collected down through a depth of ~10 cm while first clearing any leaf litter and/or debris. Replicated soil samples were homogenized and merged, split into subsamples for the various analyses, and immediately placed on ice.

### 2.2 Molecular Methods & Sequence Analysis

Both swab and soil DNA were extracted using the DNeasy PowerSoil® DNA Isolation Kit (Qiagen, Germantown, MD, USA). For bacteria, we amplified the V4 region of the 16S rRNA gene using the primer pair 515F and 806R (Apprill et al., 2015; Caporaso et al., 2011; Parada et al., 2016). For fungi, we amplified the ITS1 region using the forward primer ITS1F and reverse primer ITS2 (Gardes & Bruns, 1993; Smith & Peay, 2014; White et al., 1990). Microbial DNA was amplified in duplicate, 25-ul reactions as follows: 0.25 ul Platinum Taq (5u/ul), 0.75 ul MgCl_2_ (50mM), 0.5 ul dNTPs (10mM), 2.5 ul 10x Invitrogen Buffer, 1.0 ul of each primer (10uM), 1.0 ul BSA, and 2 ul of extracted DNA. Amplifications were performed on an Eppendorf thermocycler (Enfield, CT, USA) under the following conditions for bacteria: 94°C for 3 min, followed by 94°C for 45 s, 50°C for 1 min, 72°C for 90 s for 35 cycles, and 72°C for 10 min; and the following conditions for fungi: 94°C for 1 min, followed by 94°C for 30 sec, 58°C for 30 sec, 68°C for 30 sec for 35 cycles, and 68°C for 7 min. PCR products were purified, normalized, and pooled prior to sequencing on an Illumina MiSeq using paired-end 2 x 250-bp at GeneWiz (Brooks Life Science, South Plainfield, NJ, USA).

Demultiplexed sequences were processed using QIIME2 (Version 2019.7; Bolyen et al. 2019; Caporaso et al. 2010). For bacteria, primers and adapters were trimmed using *cutadapt* and features were truncated to remove sections of low-quality bases at the ends of each sequence as well as low-quality reads before merging via DADA2 (Callahan et al., 2016). For fungi, only the forward reads were used because of the inadequate quality of the reverse reads. Adapters, fungal primers, and low-quality bases were trimmed using *ITSXpress,* targeting the ITS1 region (Rivers et al., 2018), and *cutadapt* before further processing (Martin, 2011).

Features with less than 10 reads were removed, as they likely were sequencing artifacts, to avoid artificially inflating diversity. Reads were aligned and assigned to amplicon sequence variants (ASVs) through DADA2 and using the databases SILVA (Release 132; Quast et al., 2012) for bacteria and UNITE (Version 8.2; Abarenkov et al., 2010) for fungi. Samples were then rarified to the sampling depth of 10224 for bacteria and 5219 for fungi. Reads classified as neither bacteria nor fungi were removed prior to rarefaction.

### 2.3 Microbial Community Analysis

All post-QIIME2 analyses were performed in R version 4.0 (R Core Team 2019). Taxonomic and ASV datasets were merged using the *phyloseq* package (McMurdie & Holmes, 2014). ASVs with ambiguous phylum annotation or low prevalence ASVs (defined as < 30 reads and presence in < 5% of samples) were removed. We analyzed microbiome composition and structure on the combined salamander and soil datasets, including a description of their core microbiome (ASVs present in a minimum of 50% of samples) as well as the individual subsets of data. All *p*-values are reported as false discovery rate corrected, and visualizations were created using *ggplot2* (Wickham, 2016).

For alpha diversity, we measured ASV richness and Shannon’s diversity and performed either a Wilcox test for comparisons of season and sample type or an ANOVA for comparisons of urbanization level and site level. For beta diversity, we examined abundance-weighted analyses (Bray-Curtis) on log-transformed sequence counts. We performed comparisons of homogeneity of group dispersions (with betadispers) and PERMANOVAs using the *vegan* package (Oksanen et al., 2013) at the level of urbanization, site, and season followed by post-hoc pairwise comparisons using *pairwise.adonis* (Martinez, 2019). We used principal coordinate analysis (PCoA) to visualize beta diversity patterns. Lastly, we determined if specific ASVs were differentially abundant between sample types using DESeq2 (Love et al., 2014).

For each land-use type, we estimated the relative influence of ecological processes on the assembly of salamander bacterial and fungal communities. Following the framework developed by Stegen et al. (2013), we compared observed and null-predicted microbial communities using two beta-diversity metrics, beta nearest-taxon index (βNTI) and a modified version of the Raup-Crick index (RC_Bray_; for workflow of method, see **Figure 3a**). First, we measured phylogenetic turnover using beta mean nearest-taxon distance (βMNTD; Fine & Kembel, 2011; Webb et al., 2011) for each pairwise soil and salamander community and also quantified the degree to which observed βMNTD deviated from the null model expectation (measured in units of standard deviation, βNTI (Stegen et al., 2012)). This allowed us to estimate the influence of both homogeneous (βNTI < −2) and heterogeneous selection (βNTI > +2). Null distributions for βMNTD were generated by randomizing species in the observed data using a null model with 999 iterations that shuffles species across tips of the phylogeny (Stegen et al., 2013). For comparisons with |βNTI| < 2, we used a null model test of the Bray-Curtis taxonomic index, RC_Bray_, which modifies the Raup-Crick procedure described in Chase et al. (2011) to account for ASV relative abundances (Stegen et al., 2013). Null Bray-Curtis distributions were generated by probabilistically assembling communities 999 times while maintaining ASV richness and total read counts for each community. The difference between observed and predicted distributions were then interpreted in units of standard deviation (RC_Bray_ between −1 and +1) to partition the relative influence of stochastic processes, such as dispersal limitation and drift.

Finally, we matched ASVs from the salamander dataset with site-specific sequences of known anti-Bd bacteria and fungi in our cultured isolate dataset from salamanders at the same locations (Barnes et al., 2020) as well as with two antifungal isolate datasets (Kearns et al., 2017; Woodhams et al., 2015). To minimize overstating the putative anti-Bd function of our dataset, we only included anti-Bd information for ASVs that could be assigned to sequences from the databases at or below family level. We compared alpha and beta diversity of anti-Bd taxa across the dataset. A subset of our swab samples from salamanders were tested for Bd presence using the real-time Taqman assay developed by Boyle et al. (2004). Samples were run in duplicate alongside standards of 100, 10, 1, and 0.1 zoospore genomic equivalents (ZGEs) extracted from Bd strain JEL423, a hypervirulent strain from the global panzootic lineage *Bd*GPL. Samples were categorized as Bd-positive if both duplicates amplified before 0.1 ZGEs and all others were considered Bd-negative. In cases where only one of the duplicates returned a positive signal, the sample was run a third time and categorized based on the result of two out of three runs.

### 2.4 Soil Physicochemical Analysis

To understand how potential variation in soil physicochemical characteristics might impact microbial community composition, we conducted a range of soil analyses. Soil moisture was measured by calculating the weight difference of 5 g of wet soil after drying for 24 h at 105°C. We measured pH using an Accumet AE150 probe (ThermoFisher Scientific, Waltham, MA, USA) by mixing air dried soil with distilled water (1:2). A subset of soil samples (*n* = 45) was sent for elemental analysis (inductively coupled plasma mass spectrometry) at Bureau Veritas Mineral Laboratory (Vancouver, British Columbia, CA). Soil physicochemical variables were log-transformed and then a nested-ANOVA was performed using a mixed-effect model with level of urbanization as the fixed effect and individual site as the random effect. Post-hoc comparisons on least-mean squares were performed with a Tukey adjustment. Relationships between taxa diversity and environmental variables were determined using a canonical correspondence analysis (CCA) for both bacteria and fungi. Each CCA model was restricted to only include the environmental variables with known impacts on soil fungal or soil bacterial diversity. Model selection was performed in both directions using the ordistep function in the *vegan* package. An ANOVA was used to test the significance of the model, terms, and axes. To observe the relationship between soil-bacterial communities, soil-fungal communities, and soil physicochemical variables, pairwise-Mantel tests were run using Bray-Curtis distances and Pearson correlations on log-transformed elemental and ASV datasets.

## 3 Results

### 3.1 Overview of Microbial Diversity

After all filtering steps, our datasets generated 2.2 x 10^6^ bacterial sequences from 142 *P. cinereus* individuals and 88 soil samples and 6.0 x 10^5^ fungal sequences from 74 individuals and 88 soil samples. Our quality-filtering steps removed 4/5 samples from HH, due to low read counts and quality, thus we removed the remaining HH sample from future analyses. The full bacterial dataset was dominated by five phyla: Proteobacteria (36.2% of all bacterial reads), Acidobacteria (19.1%), Bacteroidetes (12.0%), Actinobacteria (10.6%), and Verrucomicrobia (9.0%). In total, these five phyla represented 86.9% of all reads and were common on both salamanders and in their soil environment. The 10 most abundant ASVs were from the phyla Actinobacteria (*n* = 3; Family: Micrococcaceae and Dermabacteraceae), Proteobacteria (*n* = 4; Family: Pseudomonadaceae and Xanthobacteraceae), Firmicutes (*n* = 1; Family: Bacillaceae), and Verrucomicrobia (*n* = 2; Family: Chthoniobacteraceae). When the dataset was subset by sample type, Acidobacteria was the most abundant phylum in soil with 38.6% of all reads despite representing just 8.8% of reads on salamanders in those same environments (**Figure 1a**). Following Proteobacteria, salamander skin was dominated by Actinobacteria (15.4% of all reads) whose relative abundance was 14 times higher than in the soil.

**Figure 1.**
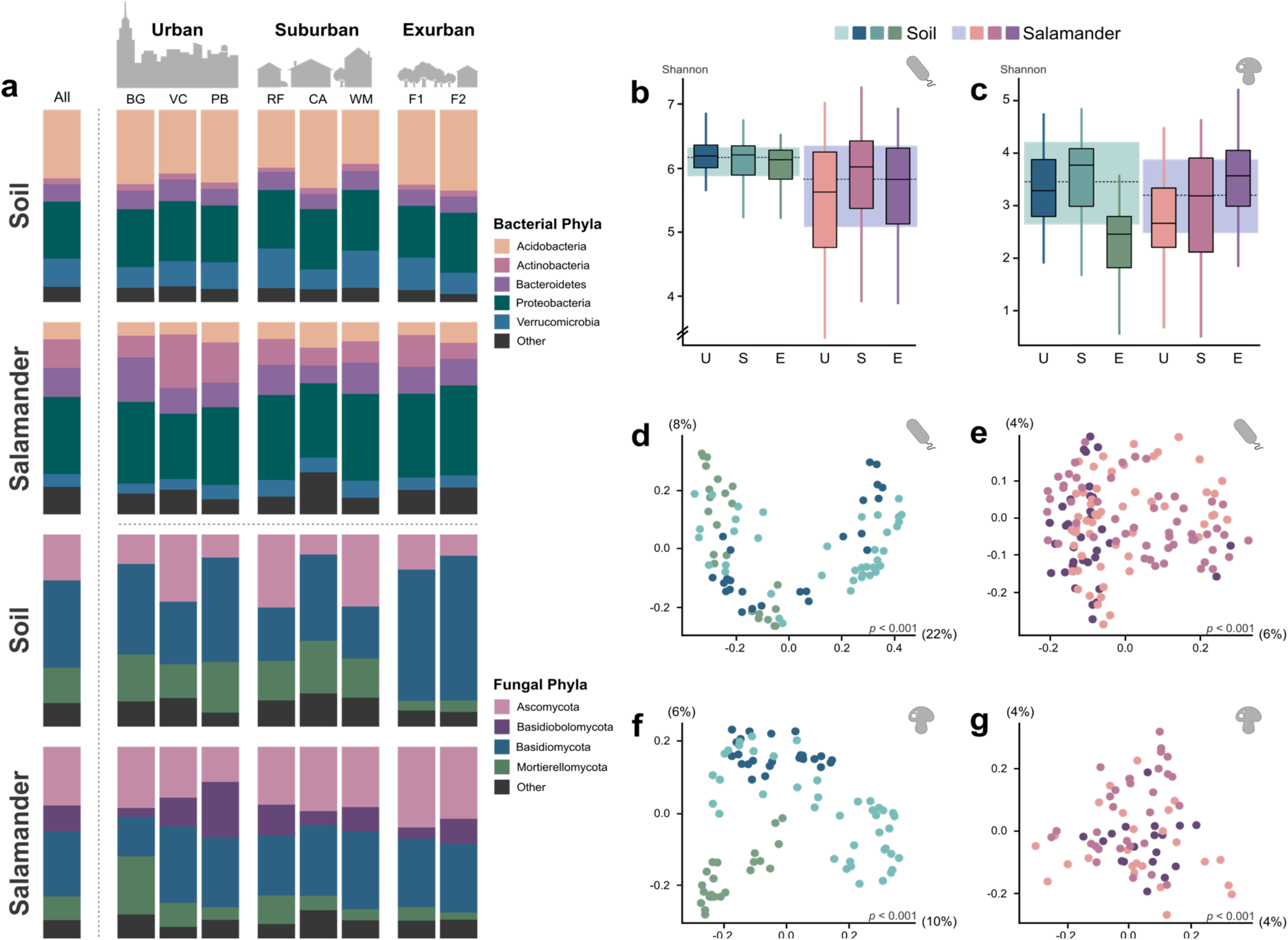
Microbial diversity of salamanders and their soil environment. a,b) Shannon diversity index of bacteria (a) and fungi (b) at each level of urbanization. Soil shown in blue-green and salamander shown in pink-purple. Boxplots show median and interquartile range with the larger box behind each level of urbanization showing the median and range for all samples of each type. c-f) PCoAs of Bray-Curtis dissimilarities for soil bacteria (c), salamander bacteria (d), soil fungi (e), and salamander fungi (f). Colors denote level of urbanization as used in alpha diversity plots (a,b). g) Relative abundance of bacterial and fungal phyla by site.

The full fungal dataset was dominated by the phylum Basidiomycota (40.6% of all fungal reads) followed by Ascomycota (27.5%), Mortierellomycota (14.7%), and Basidiobolomycota (8%; **Figure 1a**). The 10 most abundant ASVs were from the phyla Basidiomycota (*n* = 4; Family: Russulaceae and Polyporaceae), Mortierellomycota (*n* = 3; Family: Mortierellaceae), Basidiobolomycota (*n* = 2; Family: Basidiobolaceae), and Ascomycota (*n* = 1; Family: Archaeorhizomycetaceae). When subset by sample type, soil was dominated by the phyla Basidiomycota (46.1% of soil reads), Ascomycota (23.7%), and Mortierellomycota (17.6%), while the composition on salamander skin followed a similar pattern but showed a decrease in Basidiomycota (34.6% of all salamander reads) and increase in both Ascomycota (31.7%) and Basidiobolomycota (12.5%). We found the core salamander microbiome consisted of 218 bacterial taxa and 15 fungal taxa, and the core soil microbiome consisted of 172 bacterial taxa and 37 fungal taxa. Only 2 bacterial ASVs were shared by 90% of salamander samples (Family: Micrococcaceae and Xanthobacteraceae) and 5 bacterial ASVs for soil samples (Family: Chitinophagaceae, Xiphinematobacteraceae, Xanthobacteraceae, and two Acidobacteria from the genus *Bryobacter*). Finally, we found 4 fungal ASVs shared by 90% of all soil samples (Family: Piskurozymaceae, Trimorphomycetaceae, and two from Mortierellaceae).

### 3.2 Salamander and Soil Microbiomes Across Space and Time

Overall, salamander skin bacterial communities were more species-rich than those of their soil environment (salamander = 1012.9 ± 37.6, soil = 859.8 ± 29.0) but the reverse pattern was found with skin fungal communities (salamander = 80.8 ± 7.4, soil = 233.5 ± 11.3). Skin bacterial communities were less diverse than soil bacterial communities (Shannon: salamander = 5.7 ± 0.1, soil = 6.2 ± 0.04, *p <* 0.05; **Figure 1b**). We found no significant variation in the alpha diversity between skin and soil fungal communities (Shannon: salamander = 2.9 ± 0.1, soil = 3.1 ± 0.1, *p =* 0.21; **Figure 1b**). Along the land-use gradient, patterns of microbial taxonomic groups varied among richness and diversity. For bacteria, we found no significant difference in Shannon diversity for salamander or soil samples at the level of urbanization. Although, at the site level for the soil-bacterial communities, we found the suburban site RF and the urban site VC to be significantly different to the exurban site F2 (*p <* 0.05). For fungi, we found that the exurban soil communities were less diverse than the urban and suburban soil communities and the urban salamander samples were significantly different from the exurban samples (Shannon: *p <* 0.05; **Figure 1c**).

In comparisons of beta diversity, salamander microbial communities grouped separately from their soil environment in Bray-Curtis dissimilarities (bacteria: *F* = 18.2, *p* < 0.001; fungi: *F* = 12.3, *p* < 0.001). Additionally, the composition of both salamander and soil microbial communities was significantly different by both level of urbanization and site (*p* < 0.001; **Figure 1d-g**). PERMANOVA of Bray-Curtis dissimilarities showed that the site (bacteria: *R^2^* = 0.08, fungi: *R^2^* = 0.07) was the greatest predictor of differences in overall microbial community composition. This was further shown when subsetting the data for both bacteria (salamander: *R^2^* = 0.09; soil: *R^2^* = 0.17) and fungi (salamander: *R^2^* = 0.1; soil: *R^2^* = 0.18). We considered if seasonality has any effect on community composition and found it was a minimal predictor of community composition (bacteria: *R^2^* = 0.01, *p* < 0.001; fungi: *R^2^* = 0.008, *p* = 0.009).

The main differences in bacterial taxonomic composition along the land-use gradient were seen in the relative abundance of Actinobacteria, Bacteroidetes, and Verrucomicrobia. Taxa in Actinobacteria and Bacteroidetes had a non-significant but slight increase in the relative abundance in urban sites, especially BG, as compared to exurban and suburban sites. Taxa in Verrucomicrobia were significantly lower in the exurban sites as compared to sites in the other two land-use areas (*p* < 0.05). For fungi, the main differences in community composition along the land-use gradient were in the phyla Mortierellomycota and Basidiomycota. Taxa in Basidiomycota were significantly higher in exurban sites when compared to the urban and suburban sites (*p* < 0.05). Whereas exurban sites also had significantly lower Mortierellomycota taxa compared with the urban sites.

Using differential abundance analysis, we identified 1545 ASVs that were strongly associated with either sample type (536 soil-associated taxa: bacteria = 323, fungi = 213; 1009 salamander-associated taxa: bacteria = 1000, fungi = 9; *p* < 0.05). The majority of soil-associated taxa belonged to the bacterial phyla Acidobacteria and Verrucomicrobia (**Figure 2a**; top families: Acidobacteraceae (Subgroups 1, 2, and 6), Pedosphaeraceae; *p* < 0.001) and the fungal phyla Ascomycota and Basidiomycota (**Figure 2b**; top five genera in order of abundance: *Archaeorhizomyces*, *Trichoderma*, *Cenococcum*, unidentified Hydnodontaceae, *Preussia*; *p* < 0.001). The majority of skin-associated bacterial taxa belonged to the phyla Actinobacteria and Proteobacteria (top families: Micrococcaceae, Dermabacteraceae, and Sphingomonadaceae; *p* < 0.001). The majority of skin-associated fungal taxa were in Ascomycota and Basidiomycota, but from the Tremellomycetes, Microbotryomycetes, Dothideomycetes, and Leotiomycetes classes (*p* < 0.001). Of note, the ASV with the greatest log2-fold-change belonged to the genus *Basidiobolus*, the only taxa in our dataset from the phyla Basidiobolomycota.

**Figure 2.**
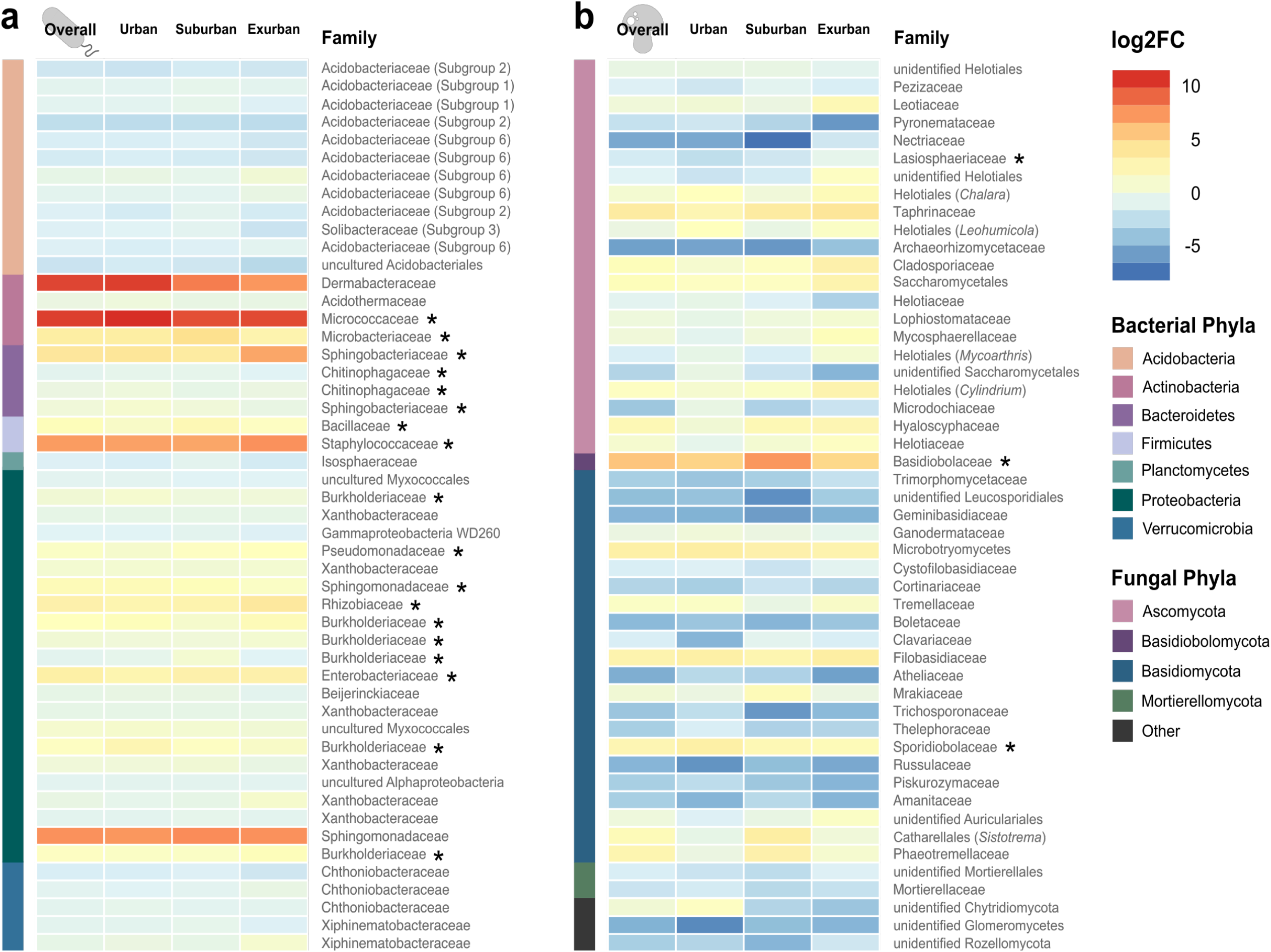
Top 50 soil-and salamander-associated microbial taxa. Heatmaps of bacterial (a) and fungal (b) ASVs that are differentially abundant between sample types. All ASVs shown were significantly more abundant in one of the two sample types (*p* < 0.05), where log2FC < 0 represents ASVs more abundant in soil and log2FC > 0 represents ASVs more abundant on salamanders. ASVs are identified by name at the family level and by color at the phylum level at the left side of each heatmap. Stars next to the family name indicate putatively anti-Bd ASVs.

To tease apart the relative importance of stochastic versus deterministic assembly mechanisms in structuring salamander microbial communities, we compared our datasets using a null-model based approach (**Figure 3b**). These models estimated that stochastic processes, such as drift and dispersal, had the largest influence on microbial assembly (76.3% ± 3.3%), especially for fungi (81.9% ± 2.1%). In contrast, deterministic processes (especially heterogeneous selection) were less pronounced (23.7% ± 3.3%) but played a larger role in bacterial assembly on salamander skin (29.3% ± 4.3%) as opposed to fungal assembly (18.1% ± 2.1%). We also found that heterogeneous selection played a much larger role in salamander skin assembly (on average, 14.4X greater for bacteria and 3.2X greater for fungi) than homogeneous selection. When we divided the dataset by level of urbanization, we found that dispersal limitation was consistently most important in governing the assembly of both urban bacteria and fungi (1.6X greater for bacteria and 1.8X greater for fungi) but that overall, undominated processes (e.g. drift) were most important in microbial community turnover (bacteria: 47.9% ± 7.4%; fungi: 76.3% ± 2.4%).

**Figure 3.**
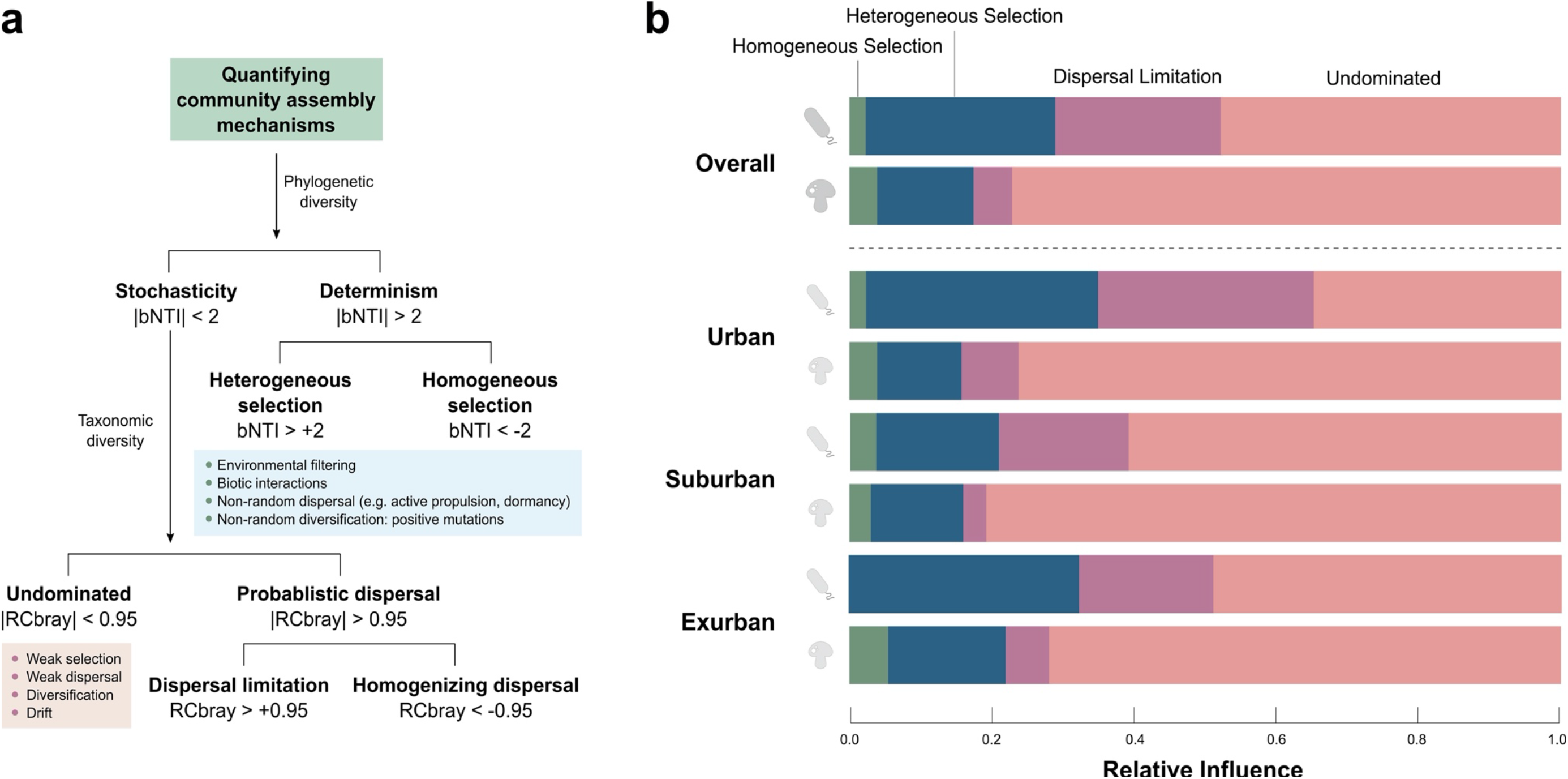
Microbial assembly mechanisms from soil to salamander skin. a) Diagram of framework developed by Stegen et al. (2013) for estimating the relative influence of stochastic and deterministic assembly processes, modified from Zhou & Ning (2017). b) Relative influence of the ecological processes shaping bacterial and fungal diversity on salamander skin across levels of urbanization.

### 3.3 Physicochemical Influence on Soil Microbial Composition

We found significant differences among five soil elements by level of urbanization: Zn, P, Ba, Al, and Rb (*p* < 0.001). Pairwise analyses showed that: a) three of these elements (Zn, Al, and Rb) were significantly different at suburban sites compared to other levels of urbanization, whereas b) Ba was significantly different between exurban and suburban sites, and c) P was significantly different at urban sites compared to other levels (*p* < 0.05). We found significant differences at the site level in these elements as well as in soil moisture, and 36 additional elements, including metals and non-metals (*p* < 0.05).

For soil bacteria, our CCA model revealed seven variables (soil moisture, canopy cover, impervious surface, K, Al, Ba, and Pb) that explained 31.4% of the variation along the land-use gradient (first axis: 11.5% explained, second axis: 4.5%, *p* < 0.001; **Figure 4a**). For soil fungi, the CCA model also revealed seven variables (distance from New York City, canopy cover, impervious surface, K, Fe, Al, and Ni) which explained 20.7% of the variation (first axis: 4% explained, second axis: 3.4%, *p* < 0.001; **Figure 4b**).

**Figure 4.**
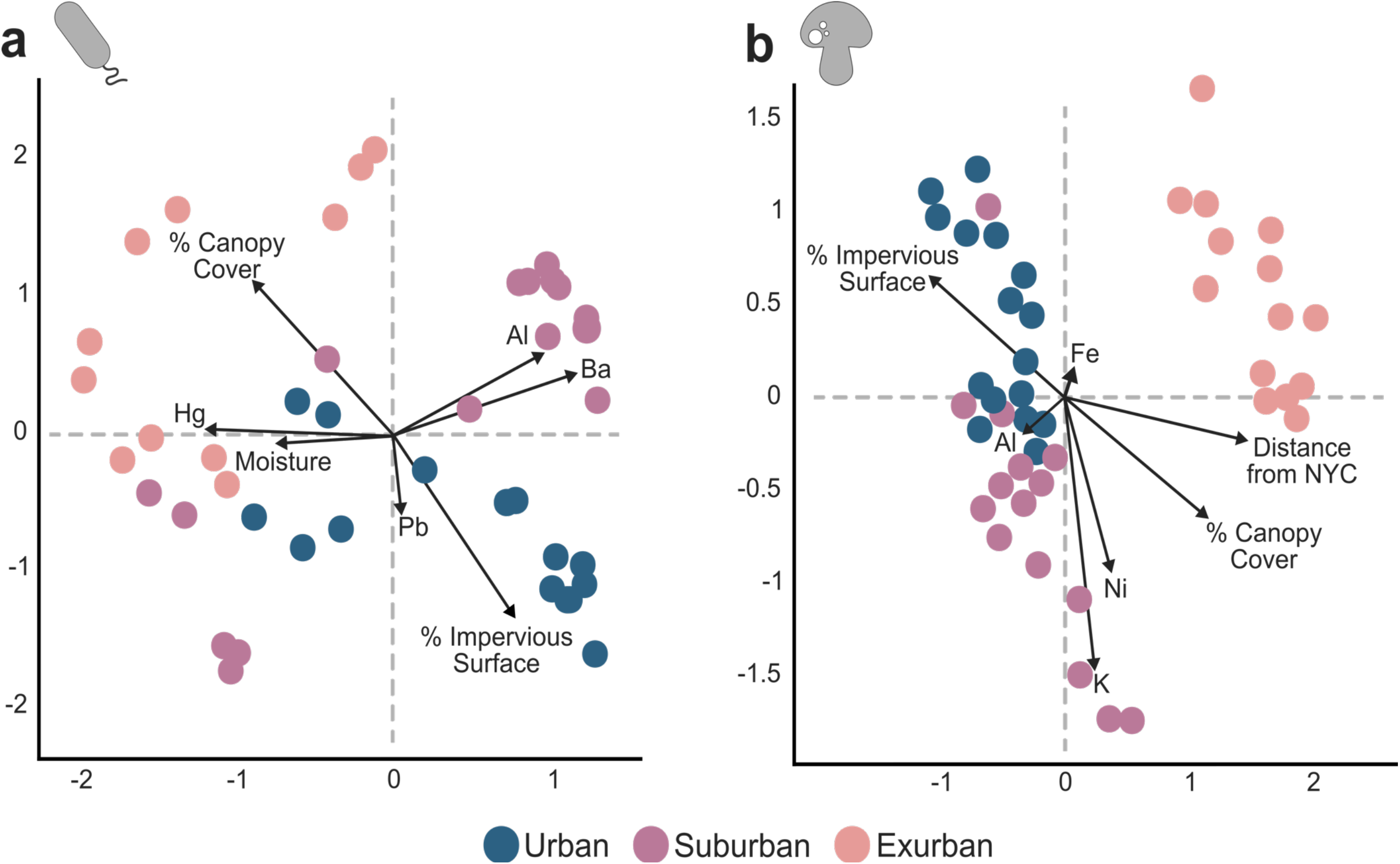
Analysis of soil physicochemical properties with soil microbial diversity. a: CCA of soil properties and bacterial diversity (*F* = 2.46, *p* < 0.001). b: CCA of soil properties and fungal diversity (*F* = 1.38, *p* < 0.001). Dots represent individual soil samples and are colored by level of urbanization.

We next compared the fungal and bacterial communities to soil environmental variables (all Mantel correlation coefficients reported in **Table 1**; all comparisons were considered significant if *p* < 0.01). Overall, soil bacteria were most correlated with percent impervious surface (*r* = 0.25), percent canopy cover (*r* = 0.24), and Al concentration (*r* = 0.19). Additionally, Acidobacteria and Proteobacteria were most correlated with Pb (*r* = 0.24; *r* = 0.19) and Al (*r* = 0.16; *r* = 0.21), Actinobacteria with Al (*r* = 0.18) and Cu (*r* = 0.12), and Bacteroidetes with Pb (*r* = 0.17) and Zn (*r* = 0.19). Similar to soil bacteria, fungi were most correlated with percent impervious surface (*r* = 0.45), percent canopy cover (*r* = 0.26), and Al concentration (*r* = 0.27). All three dominant fungal phyla also correlated with Zn (Basidiomycota: *r* = 0.13, Ascomycota: *r* = 0.19, Mortierellomycota: *r* = 0.17) and Na (Basidiomycota: *r* = 0.22, Ascomycota: *r* = 0.22, Mortierellomycota: *r* = 0.28).

**Table 1.**
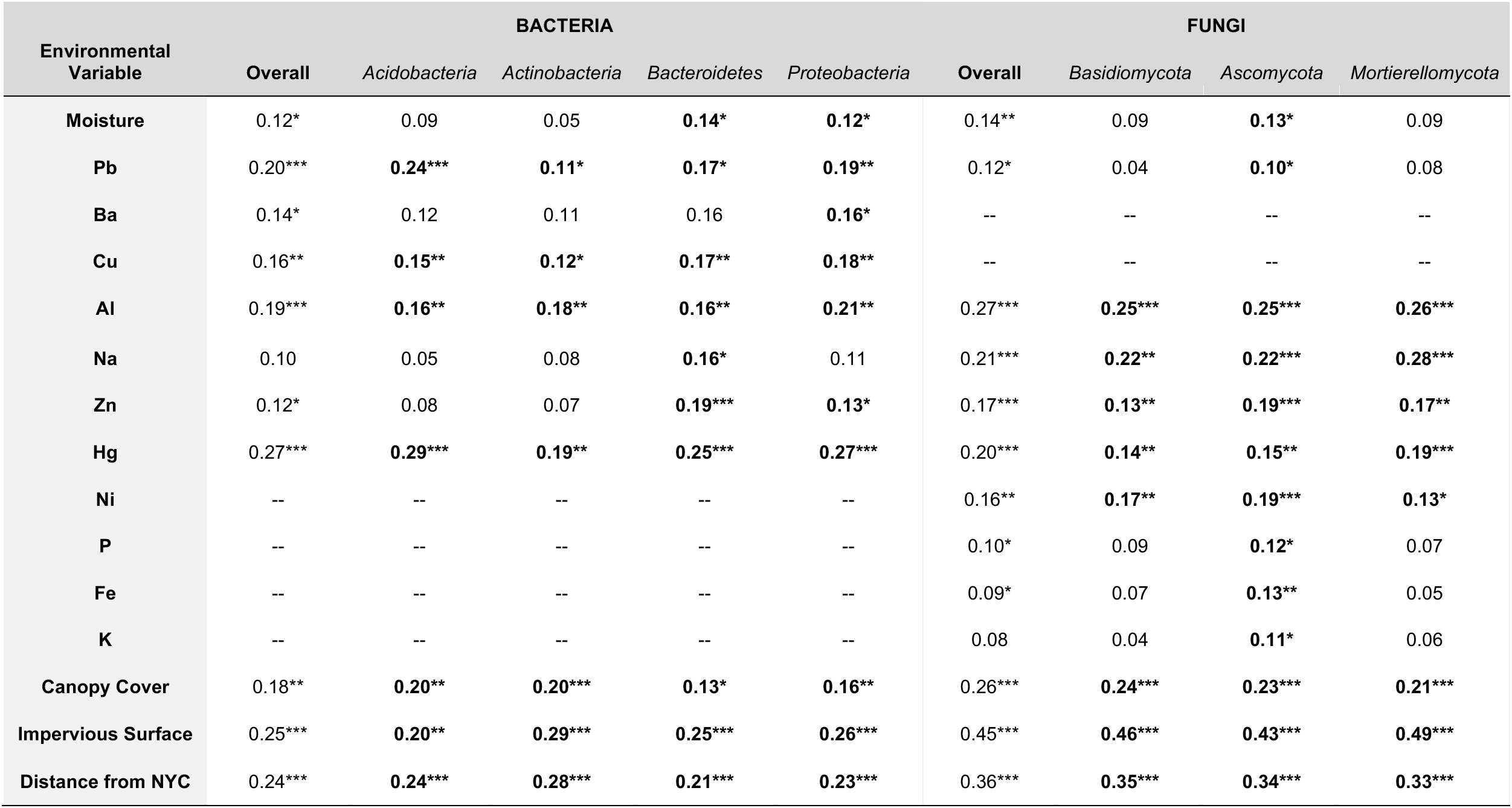
Mantel correlations between soil-microbial diversity and physicochemical variables. Both beta diversity (Bray-Curtis dissimilarities) and environmental matrices were log-transformed prior to analysis. Correlations were identified overall for the bacterial and fungal communities as well as for dominant phyla. Significant correlations identified with one (*p* < 0.05), two (*p* < 0.01), or three (*p* < 0.001) stars and shown in bold for individual phyla. Dashes identify variables not found to be important to the CCA models of those groups.

### 3.4 Distribution of Anti-Bd microbes

Of the 82 salamander samples tested for Bd, twenty were positive (> 0.1 ZGE): 12% of urban, 33% of suburban, and 28% of exurban samples, but all showed very low zoospore abundance (mean = 1.1 ± 0.4 ZGE). Interestingly, we found no significant difference in the microbial communities of salamanders that tested positive or negative for Bd (*F* = 0.82, *p* = 0.91), but the composition of putatively anti-Bd microbes varied over space and time (**Figure 5**). Overall, we found 1402 putatively anti-Bd bacterial ASVs and 248 anti-Bd fungal ASVs. Our combined anti-fungal database identified 32 microbial genera as non-inhibitory or facilitatory but lacked information on a large portion of the bacterial and fungal taxa observed. Anti-Bd ASVs made up ~28% of all bacterial ASVs and ~2.5% of all fungal ASVs. At the individual salamander level, samples varied in anti-Bd ASV relative abundance from <1% to 68% of sequences per sample.

**Figure 5.**
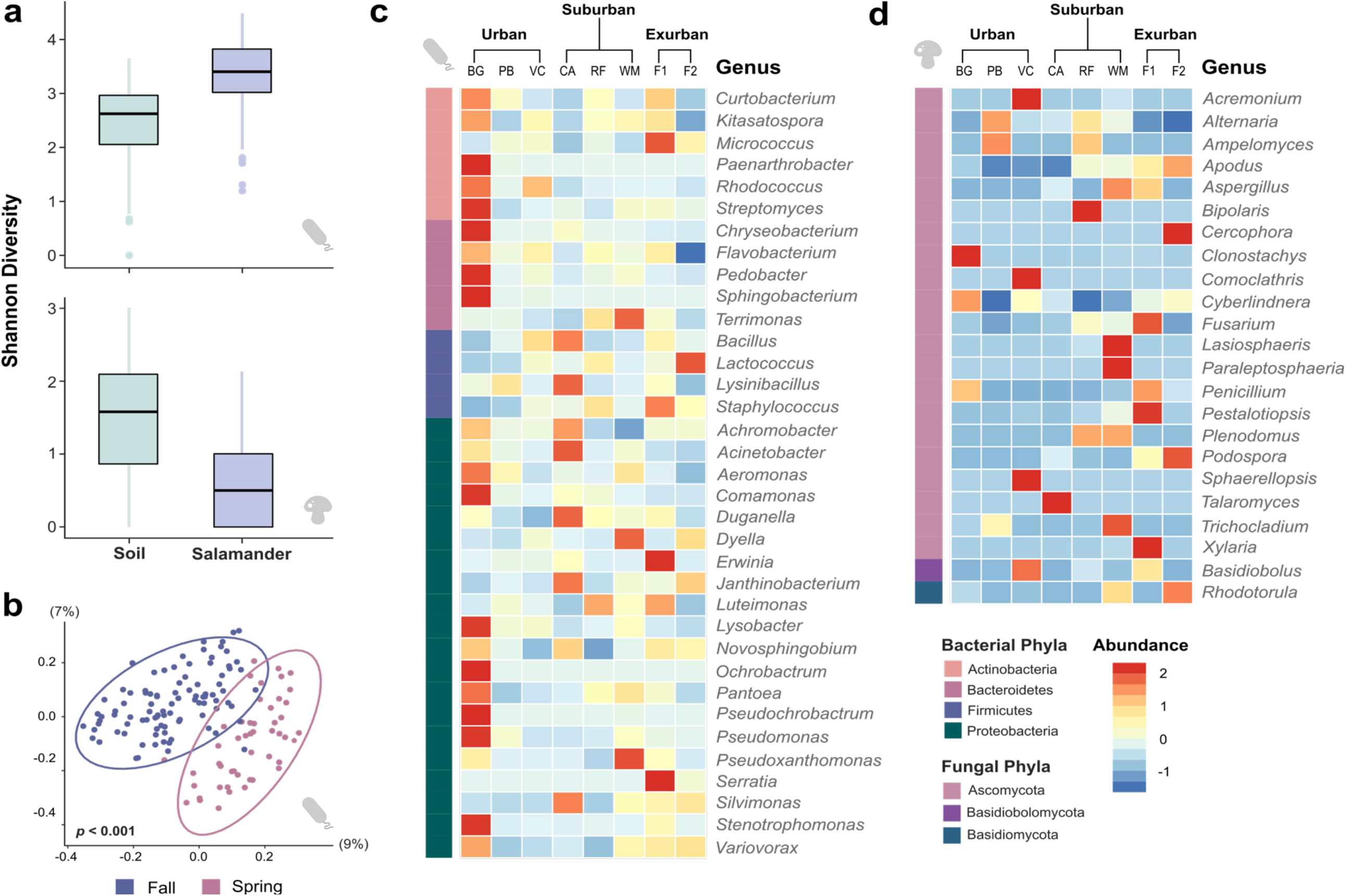
Diversity of putatively anti-Bd bacteria and fungi on *Plethodon cinereus* skin. a: Shannon diversity in soil and on salamander hosts. b) PCoA of Bray-Curtis dissimilarities between anti-Bd salamander bacterial communities in Fall and Spring. Heatmap of z-score standardized abundances of putatively anti-Bd bacterial (c) and fungal (d) genera found on salamanders across the gradient of land-use change. Site-level Z-score abundances < 0 denote ASVs belonging to that genus are less than the mean abundance for that genus. Corresponding phylum of each taxa is identified on the left side of each heatmap.

Overall, both anti-Bd bacteria and fungi represented a larger proportion of of the salamander microbiome than the soil microbiome (salamander: 14% of reads for bacteria and 1.5% for fungi; soil: 2% of reads for bacteria and 0.9% for fungi) regardless of level of urbanization. Additionally, salamanders hosted a more diverse community of anti-Bd bacteria, but a less diverse community of anti-Bd fungi than the soil (*p* < 0.001; **Figure 5a**). In particular, the core salamander skin anti-Bd microbiome consisted of 19 bacterial taxa and just 2 fungal taxa, which represented < 1% of all putatively Bd-inhibitory ASVs observed. When we analyzed the dataset across time, we found that salamanders sampled in the fall had more species-rich anti-Bd bacterial communities than in the spring (*p* = 0.003) leading to separation of these communities in our ordination of Bray-Curtis dissimilarities (*F* = 8.16, *p* < 0.001; **Figure 5b**).

However, we found no significant difference between the anti-Bd fungal communities of salamanders in Fall or Spring. While anti-Bd bacterial beta diversity appeared significantly different by site (*F* = 2.7801, *p* < 0.001), significant differences in composition were mainly driven by one site, BG (*F* = 2.3022, *p* = 0.03; all sites compared to BG: *p* < 0.05; **Figure 5c**). Alternatively, anti-Bd salamander fungal communities did vary significantly by site (*p* < 0.05; **Figure 5d**). By looking at the abundances of individual taxa across the gradient, we found that the bacteria *Serratia* and the fungi *Apodus* decreased with increasing urbanization (*p* < 0.001). Additionally, the well-known anti-Bd bacterial genus, *Janthinobacterium*, appeared in relatively very low abundances in all urban sites while other well-known anti-Bd genera *Flavobacterium* and *Lysobacter* appeared in relatively high abundances in urban sites.

## 4 Discussion

This work examines both the local and regional factors influencing the assembly of the skin microbiome in the amphibian species, *Plethodon cinereus*. By surveying both the bacterial and fungal communities across a gradient of land-use change, we were able to identify how variation in the environment can create high dissimilarity across individuals of a single host species. In particular, we found a significant influence of a number of physicochemical factors on the soil microbial reservoir that supplies colonists to amphibian skin. This coincided with variation in both the bacterial and fungal communities of salamanders found at these sites. We believe our findings provide valuable insight into the exogenous factors often confounded by host species identity, a well-supported predictor of microbial composition (Ellison et al., 2018; Jani & Briggs, 2018; Muletz-Wolz et al., 2018). Additionally, this study reveals the important roles that dispersal, drift, and heterogeneous selection play in driving dissimilarity between individual hosts.

### 4.1 Land-use change alters microbial composition in the environmental reservoir

We found that soil bacterial and fungal communities differed in their response to land-use change. Despite ongoing habitat degradation, it seems that the bacterial phyla Acidobacteria, Proteobacteria, and Verrucomicrobia remain highly abundant. These phyla are common to soil environments across the globe where they are associated with carbon usage and nitrogen assimilation (Janssen, 2006; Kielak et al., 2016; Sait et al., 2002). However, the relative abundance of these phyla is known to vary with certain environmental conditions. When we compared beta diversity across the gradient, we found individual ASVs within each bacterial community were highly heterogeneous. Additionally, each major bacterial phylum was correlated with a diverse set of soil physicochemical characteristics that are difficult to disentangle from site-specific effects and regional urbanization. Since our samples came from woodlands across a relatively small spatial scale and our sites did not significantly vary in pH (Dequiedt et al., 2011; Fierer & Jackson, 2006; Hansel et al., 2008), this may explain why we didn’t see significant differences in bacterial diversity across the gradient.

In contrast to bacteria, the relative abundance of the common soil fungal phyla, Basidiomycota and Mortierellomycota, appeared sensitive to land-use change. These findings are consistent with other urban-focused studies that suggest that urbanization might drive homogenization of soil fungal communities towards increased abundances of Mortierellomycota and Ascomycota (Abrego et al., 2020; Becker & Lewis, 2012; Epp Schmidt et al., 2017; Jurburg et al., 2020; Karpati et al., 2011; van Geel et al., 2018). In turn, this could lead to decreases in functionally important taxa within Basidiomycota (Abrego et al., 2020; Epp Schmidt et al., 2017), which may be dispersal limited or are not as resilient to the impacts of land-use change. There remains the possibility that these communities have gone through an irreversible change in composition following historical land-use changes (Clemmensen et al., 2015; Cline et al., 2015; Dupouey et al., 2002). For instance, saprotrophic taxa within Ascomycota and Mortierellomycota, known early colonizers, could inhibit the colonization of late arriving cord forming mycorrhizal taxa within Basidiomycota.

### 4.2 Salamander microbiomes are distinct, but connected to their soil environment

Salamander microbial communities showed a decreased relative abundance of Acidobacteria and an increase in Actinobacteria and Proteobacteria taxa relative to the soil. Differences between soil and salamander microbiomes likely result from a change in environmental factors which favor or select for certain microbial taxa over others (Medina et al., 2019; Rebollar et al., 2016; Walke et al., 2014). These shifts in abundance could be driven by the presence of skin antimicrobial peptides, skin-shedding, diet, and the host’s immune system which act selectively to exclude Acidobacteria (Antwis et al., 2014; Huang et al., 2016; Meyer et al., 2012; Muletz-Wolz et al., 2018). In fact, we found that many of the taxa in the salamander microbiome were absent from the soil microbiome, suggesting that amphibians either acquire their microbiome from multiple sources (e.g. conspecifics) or that these taxa were present at very low abundances in soil (Bird et al., 2018). We also found an increase in the abundance of ASVs known to produce antibiotic and antifungal compounds, particularly from the phyla Actinobacteria and Proteobacteria, on salamander skin (Barka et al., 2016; Bradley & Pollard, 2017; Cosseau et al., 2016; Ludwig et al., 2015; Rizzatti et al., 2017). As host symbionts, these ASVs likely represent the first line of amphibian immune defense to skin pathogens and their presence across space and time is promising with regard to developing microbial treatments for disease.

Our results show that salamander fungal communities are dominated by Ascomycota and Basidiomycota taxa. These results align with the few studies conducted on other amphibian species, suggesting that taxa from these phyla seem to be common regardless of amphibian species or location (Kearns et al., 2017; Medina et al., 2019; Rebollar et al., 2016). Of particular note, the fungal communities of the *P. cinereus* sampled here showed an increased relative abundance of the genus *Basidiobolus* than the soil environment. *Basidiobolus ranarum* is a common amphibian gut microbe which likely is the primary source of this taxa to the skin microbiome via fecal contamination (Green & Munths, 2005; Muths et al., 2003). However, due to the nature of our sampling and the size of *P. cinereus* individuals, we were unable to associate the presence of *B. ranarum* with proximity to the mouth or cloaca. Similar to bacteria, many of the fungal taxa identified here produce antibacterial and antifungal metabolites so these host-associated communities could be an important facet to host defense and skin function (Kearns et al., 2017; Rebollar et al., 2016).

### 4.3 Stochastic processes drive high dissimilarity between salamander hosts

In addition to the general compositional differences seen on salamander hosts between sites, our assembly model identified high dissimilarity between individuals despite no significant difference in alpha diversity. This suggests that the composition of their microbiome is shaped by more than just the size and diversity of the regional species pool (Chase, 2010; Kraft et al., 2011; Myers et al., 2013; Tucker et al., 2016). Our model revealed that heterogeneous selection played a much larger role in salamander skin assembly (on average, 14X greater for bacteria and 3X greater for fungi) than homogeneous selection. The increased importance of heterogeneous selection suggests that host species is not the only selective mechanism regulating amphibian microbiome variation. One explanation for this could be that individuals may have different immune responses to environmental stressors that change skin conditions, indirectly affecting the microbial species that establish (Rollins-Smith et al., 2011). These communities are then likely shaped by microbial interactions between colonizing and established skin microbes.

However, even after taking into account the environment, season, and Bd-presence, much of the variation in salamander bacterial and fungal communities was left unexplained. This could be related to the combined effect of phylogenetic underdispersion of microbes which can drive strong priority effects during microbiome assembly. Such events likely create alternative states of functionally redundant microbiomes among individuals despite high dissimilarity at the level of ASVs (Aguirre de Cárcer, 2019). Most importantly, the high dissimilarity found between individuals from the same site and season combined with the dominating influence of stochastic processes in our model, suggests that individual salamander microbiomes are largely shaped by assembly processes such as dispersal and drift. The influence of these processes was expected at the regional scale but was surprisingly high at the local scale. This adds to the growing literature conveying the importance of these processes in shaping microbial communities especially under land-use change (Adair & Douglas, 2017; Dumbrell et al., 2010; Martiny et al., 2011; Ofiţeru et al., 2010).

### 4.4 Host microbiome assembly under disease

Stochastic and deterministic assembly mechanisms likely play an important role in determining host susceptibility to diseases such as Bd infection. Interactions between Bd and bacteria have been documented as strong drivers of amphibian microbiome composition (Becker et al., 2019; Becker & Zamudio, 2011; Flechas et al., 2018; Jani & Briggs, 2014).

Although 25% of the individuals that we tested for Bd were positive, all Bd-positive individuals carried low pathogen loads, which may explain why, unlike in other studies, we found no significant difference in the communities of Bd-positive and Bd-negative salamanders (Bletz et al. 2017; Familiar-López et al., 2017). This also could be characteristic of this species as they are generally believed to not be at risk of extinction due to Bd (Becker & Harris, 2010; Harris et al., 2006). In line with this, we found that anti-Bd ASVs increased in abundance on salamanders as compared to their soil environment, but that composition of anti-Bd ASVs varied with site and land-use change. Thus, land-use change might impact salamander microbiomes both directly by inducing immune responses on their skin and indirectly by altering physicochemical properties of the soil which supplies colonists to amphibian skin. With increasing land-use change and expansion of Bd (Scheele et al., 2019b), many amphibian species may experience habitat shifts, altering the microorganisms they come in contact with (Cohen et al., 2019; Scheele et al., 2019a; Varela et al., 2018), making our approach all the more relevant to achieving conservation aims.

### 4.5 Conclusion

In sum, this study offers new insights into the role of both the environment and disease in regulating composition and structure of microbial species pools and local communities. Through in-depth exploration of *Plethodon cinereus* microbiomes across a gradient of land-use, we connect habitat-filtering mechanisms in the environment to small but significant effects on individual host microbiome assembly. Additionally, we identify an understudied but important role of stochastic and deterministic assembly mechanisms in driving alternative states of host microbial diversity. We believe these findings will improve our understanding of how beneficial microbiome function is created and maintained aiding in conservation efforts of susceptible taxa.

## Declarations

### Ethics Approval

All procedures involving animals was granted by the Fordham University Institutional Animal Care and Use Committee (IACUC #: JL-17-01) and the following agencies: New York State Department of Environmental Conservation (License #: 2203), New York State Office of Parks, Recreation, and Historic Preservation (Permit #: 2017-MP15-010), and the New York City Department of Parks and Recreation Natural Resources Group (Years: 2017-2019).

## Acknowledgments

We thank the managers at all of our research sites for access to soil and salamander sampling, including Rockefeller State Park Preserve, Fahnestock State Park, the New York Botanical Gardens, the Westmoreland Sanctuary, the Louis Calder Center, Van Cortlandt Park, and Pelham Bay Park. We thank Kiara Taveras, Julia Piccirillo-Stosser, and Sabrina Piccirillo-Stosser for their assistance with aspects of the soil physicochemical analysis. We also thank Drs. Katie Schneider-Paolantonio, Steven Franks, John Wehr, and Patricio Meneses for their feedback on earlier drafts of this manuscript.

---

**Table S1.**
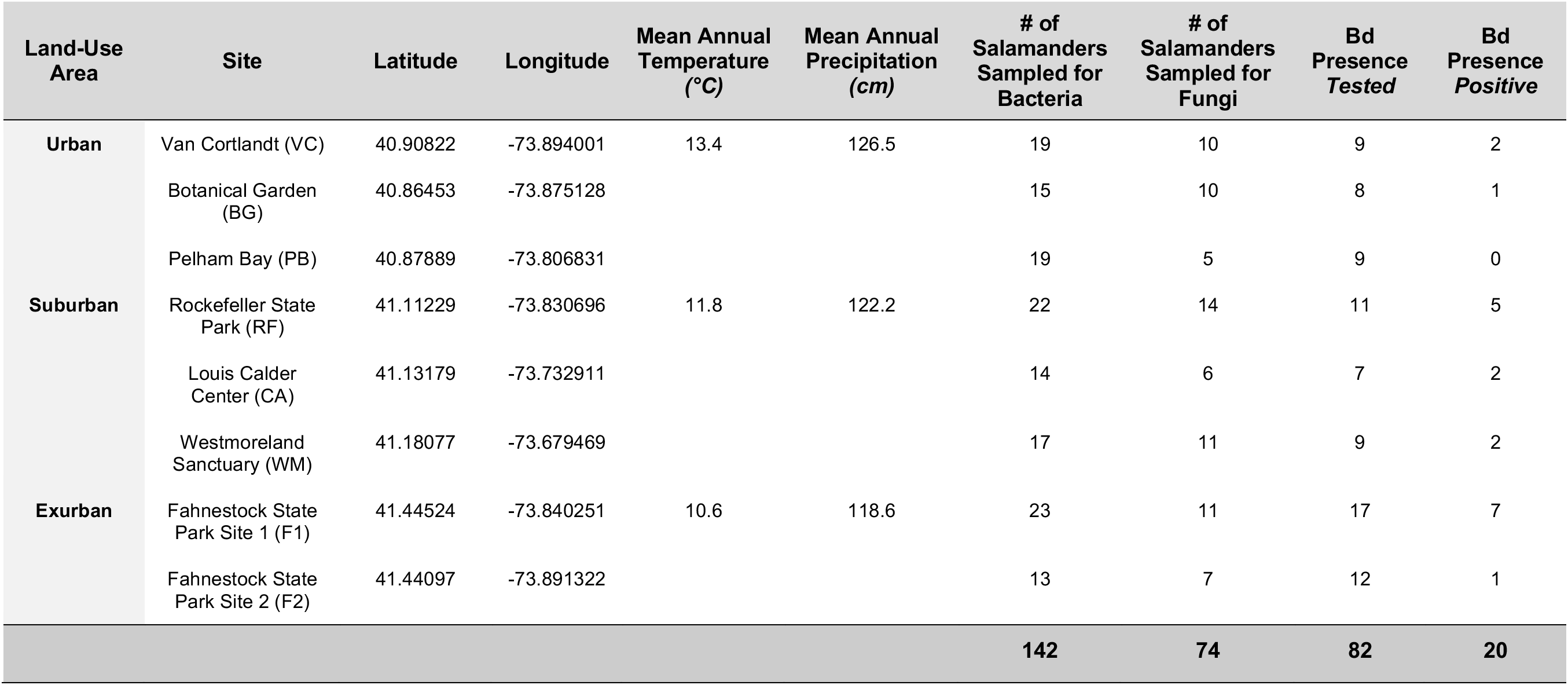
Summary of study sites and salamander sample information along land-use gradient. A subset of 82 salamanders were tested for Bd presence (ZSEminimum = 0.1). Climatic environmental variables obtained from annual temperature and precipitation data from the National Oceanic and Atmospheric Administration data repositories.

